# A Vma13-targeted thiadiazole-succinimide disrupts V-ATPase dependent arginine-polyamine homeostasis to block *Candida albicans* escape from macrophages

**DOI:** 10.64898/2026.10.29.762984

**Authors:** Xinyu Fang, Bingchen Jiang, Wanqian Li, Xiaojing Xu, Liping Xu, Xiao Zhou, Wenqing Shi, Yuanying Jiang, Hui Lu

## Abstract

Invasive candidiasis caused by *Candida albicans* is a life-threatening nosocomial infection with limited therapies and high morbidity and mortality among immunocompromised patients. Within host phagocytes, *C. albicans* catabolizes arginine to produce ammonia and fuels polyamine biosynthesis, which correlates with phagosomal alkalinization and yeast-to-hypha morphogenesis, facilitating macrophage escape and systemic dissemination. However, new antifungal strategies blocking this fungal escape remain scarce. Here, we characterize the thiadiazole-succinimide derivative ETSI, an arginine-dependent antifungal small molecule with minimal direct growth inhibition under arginine-free conditions yet robust anti-*C. albicans* activity in arginine-rich microenvironments mimicking the macrophage intracellular niche. Mechanistically, ETSI targets the vacuolar Vma13 to inhibit vacuolar H-ATPase (V-ATPase), dissipating vacuolar proton (H^+^) gradients and leading to cytoplasmic H^+^ retention. The accumulated cytoplasmic H^+^ traps intracellular ammonia and blocks its external release, while impaired vacuolar metabolite sequestration further triggers cytotoxic accumulation of arginine and polyamine metabolites. This dual metabolic perturbation suppresses phagosomal alkalinization, retains *C. albicans* in yeast form and blocks macrophage escape. ETSI is the first small molecule blocking *C. albicans* escape via disrupted Vma13/V-ATPase dependent arginine-polyamine homeostasis. This work identifies an intracellular metabolic vulnerability and establishes a niche-targeted antifungal strategy distinct from classic growth-suppressive drugs for treating invasive candidiasis.

## Introduction

Invasive candidiasis, primarily caused by the opportunistic pathogen *Candida albicans*, is a life-threatening nosocomial infection with high morbidity and mortality among immunocompromised patients worldwide[1, 2]. Current clinical antifungal arsenals are limited to three major classes, including azoles, echinocandins and polyenes, all of which primarily target fungal growth and viability[3, 4]. Worryingly, the widespread emergence of antifungal resistance, intrinsic host toxicity, and poor intracellular penetration severely compromise the efficacy of conventional antifungal agents[5, 6]. Furthermore, these growth-inhibitory drugs fail to interfere with the adaptive metabolic mechanisms that enable *C. albicans* survival and immune escape within host cells, highlighting an urgent unmet need for innovative antifungal strategies targeting fungal in vivo fitness and immune evasion[7–9].

Macrophages are the primary innate immune cells that eliminate invading *C. albicans* via phagocytosis and lysosomal acidification of phagosomes[10, 11]. However, *C. albicans* has evolved a sophisticated arginine dependent metabolic programme to neutralize phagosomal acidity, undergo yeast-to-hypha morphogenesis, and escape macrophage clearance, driving persistent intracellular infection and systemic dissemination[10, 12]. Upon phagocytosis, *C. albicans* cells import host derived arginine through Stp2 regulated amino acid permeases[13, 14]. Internalized arginine is sequentially catabolized by arginase Car1 to generate ornithine and urea[15, 16], and urea is further decomposed by urea amidolyase Dur1,2 into ammonia (NH₃) and carbon dioxide (CO_2_)[15, 16]. Vacuolar proton (H^+^)-ATPase (V-ATPase) is assembled from a cytosolic ATP-hydrolyzing V_1_ domain and a membrane-embedded V_0_ domain, which cooperatively pump protons across the vacuolar membrane to establish a stable proton electrochemical gradient. Under physiological cytoplasmic conditions maintained by V-ATPase, which pumps cytoplasmic protons into the vacuole to sustain cytoplasmic pH homeostasis[17, 18], uncharged NH_3_ readily diffuses across the fungal plasma membrane[19]. The exported ammonia substantially alkalinizes the normally acidic phagosomal lumen, which is a prerequisite for *C. albicans* intracellular survival and subsequent immune escape[13, 20]. When cytoplasmic H^+^ accumulates, NH_3_ is protonated into charged ammonium ions (NH_4_^+^) that cannot efficiently cross lipid bilayers, thereby blocking NH_3_ efflux[19]. Nevertheless, small-molecule strategies specifically disrupting this NH_3_ efflux-dependent immune escape pathway in *C. albicans* remain largely unexplored.

Beyond fuelling the ammonia-generating escape cascade, arginine also feeds a parallel biosynthetic branch for polyamine production, which is physiologically essential for fungal homeostasis and morphogenesis. In this pathway, Car1 derived ornithine is converted to putrescine by ornithine decarboxylase Spe1; putrescine is further sequentially transformed into spermidine and spermine via the catalysis of Spe3 and Spe4, respectively[15, 21, 22]. Additionally, Spe2 mediates the generation of decarboxylated S-adenosylmethionine (dcSAM) to provide critical aminopropyl donors for polyamine chain elongation[21]. While basal levels of intracellular polyamines support normal fungal physiology, excessive accumulation of polyamines causes prominent cytotoxic stress and impairs fungal viability[23]. V-ATPase acts as a central upstream regulator to maintain arginine-polyamine metabolic homeostasis in *C. albicans*. The proton electrochemical gradient generated by V-ATPase supplies the driving force for vacuolar sequestration and compartmentalization of polyamines and related metabolic intermediate metabolites, preventing toxic polyamine build-up within the cytosol[24, 25]. Given the strict dependence of vacuolar polyamine sequestration on H^+^ gradient driven transport, V-ATPase dysfunction is predicted to collapse vacuolar compartmentalization and induce intracellular accumulation of arginine and polyamine intermediates. To date, no small-molecule compounds have been reported to target this V-ATPase dependent metabolic regulatory axis to suppress *C. albicans* immune evasion.

Thiadiazole-succinimide scaffolds exhibit diverse biological activities in medicinal chemistry [26], yet their antifungal potential remains largely underexplored. In this work, we functionally characterize a thiadiazole-succinimide derivative ETSI (1-(4-ethoxyphenyl)-3-[(5-methyl-1,3,4-thiadiazol-2-yl)sulfanyl]pyrrolidine-2,5-dione) as a unique arginine dependent antifungal agent, whose chemical structure is shown in Figure 1a. ETSI displays negligible direct growth inhibitory activity under arginine free conditions yet it exerts robust anti-*C. albicans* activity within arginine rich microenvironments that mimic the macrophage intracellular niche. Mechanistically, ETSI targets Vma13 and efficiently suppresses V-ATPase H^+^ pumping activity, causing cytoplasmic H^+^ accumulation. This H^+^ overload blocks efflux and collapses the V-ATPase dependent gradient required for vacuolar polyamine compartmentalization, leading to toxic polyamine buildup. Combined, these metabolic perturbations inhibit yeast-to-hypha morphogenesis and prevent *C. albicans* escape from macrophages. Collectively, these findings establish a previously unrecognized dual mechanism linking V-ATPase mediated H^+^ balance to fungal ammonia efflux and polyamine toxicity, providing a novel niche-targeted paradigm for intervening in invasive candidiasis.

**Figure 1.**
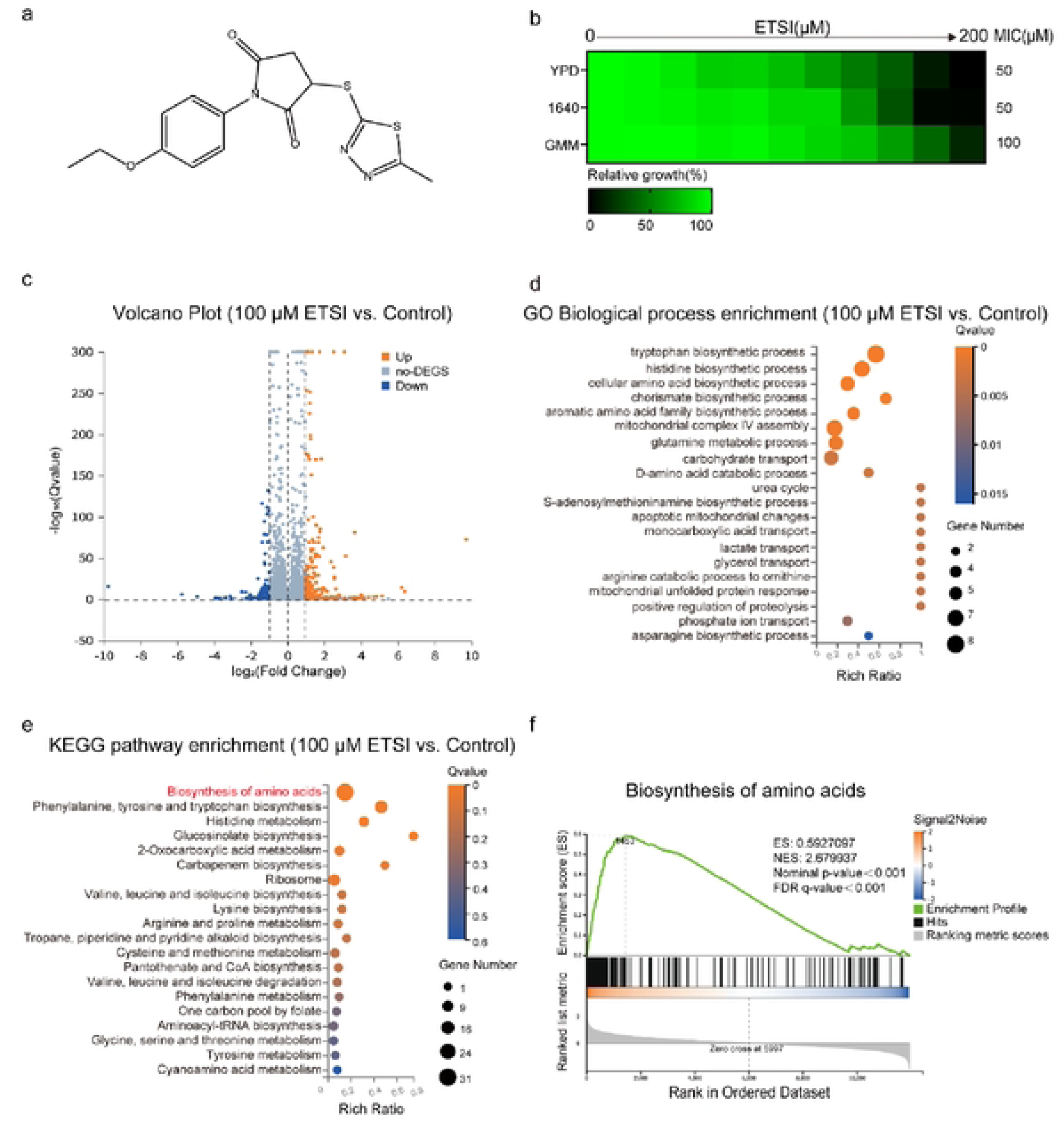
Compound ETSI induces transcription of amino acid biosynthesis pathways in *C. albicans*, **a.** Structure of compound ETSI. **b.** Determination of the MIC of compound ETSI against *Candida albicans* in YPD, RPMI 1640, and GMM media. The MIC was defined as the lowest concentration of compound ETSI that resulted in more than 50% growth inhibition, c. Volcano plot of differentially expressed genes in *C. albicans* following treatment with 100pM (2 × MIC) compound ETSI relative to the untreated control. Differentially expressed genes (DEGs) were defined as those with | log_2_(fold change) i > 1 and Q < 0.001. Upregulatcd, down regulated, and non-differentially expressed genes (non-DEGs) are shown in orange, blue, and gray, respectively, d. GO Biological Process enrichment analysis of *C. albicans* samples treated with lOOμM (2 × MIC) compound ETSI. The results indicated that compound ETSI primarily affected multiple biological processes related to amino acid biosynthesis, e. KEGG pathway analysis revealed that lOOμM (2 × MIC) compound ETSI primarily affected biological processes associated with amino acid biosynthesis, f. Gene set enrichment analysis (GSEA) demonstrated an upregulated trend in amino acid biosynthesis pathways. The enrichment score (ES) indicated that genes involved in amino acid biosynthesis pathways were enriched and upregulated following compound ETSI treatment.

## Results

### Compound ETSI upregulates amino acid biosynthesis pathways at the transcriptional level in *C. albicans*

Previous studies have documented that nitrogen and carbon source availability modulates the antifungal activity of compounds[27–29]. Yeast extract peptone dextrose (YPD) medium is nitrogen-replete and supplemented with 2% (w/v) glucose, whereas glucose minimal medium (GMM) (containing 0.67% (w/v) YNB, 0.5% (w/v) (NH₄)₂SO₄, and 2% (w/v) glucose) used in this study lacks exogenous amino acid supplements but contains identical glucose concentrations. RPMI 1640 medium recapitulates key nutritional features of human blood and contains only 0.2% (w/v) glucose. We first determined minimum inhibitory concentration (MIC) values of ETSI against *C. albicans* cultured in YPD, GMM, and RPMI 1640 media. ETSI exhibited an MIC of 50 μM in YPD and RPMI 1640 media, whereas the MIC increased to 100 μM in GMM medium (Fig. 1b). These observations indicate that the antifungal activity of ETSI is modulated by nitrogen nutrient availability, and that higher environmental nitrogen supports stronger antifungal potency of ETSI.

To dissect the mechanisms of action of ETSI, we then carried out transcriptomic profiling of *C. albicans. C. albicans* cells were treated with ETSI at 25 μM (1/2×MIC), 50 μM (1× MIC), and 100 μM (2× MIC), with vehicle-treated cells serving as controls. Compared with the vehicle control, 55 genes were up-regulated and 19 genes were down-regulated upon 25 μM ETSI treatment (Fig. S1a); 87 up-regulated and 30 down-regulated genes at 50 μM (Fig. S1b); and 172 up-regulated and 54 down-regulated genes at 100 μM (Fig. 1c) (Table S1). Gene Ontology (GO) Biological Process (BP) enrichment revealed prominent perturbation of amino acid related metabolic and biosynthetic programmes, including tryptophan, histidine, and aromatic amino acid biosynthesis (Fig. 1d). GO Molecular Function (MF) enrichment further highlighted multiple enzymatic activities linked to amino acid turnover, such as anthranilate synthase, branched-chain amino acid transaminase, S-adenosylmethionine decarboxylase, and arginase activities, pointing toward altered arginine-polyamine metabolism upon ETSI exposure (Fig.S1c). GO Cellular Component (CC) analysis showed that differentially expressed gene products were enriched in the cytosol, cell wall, extracellular region, and mitochondrial respiratory chain, as well as vacuole associated membrane components (Fig.S1d). Consistent with GO results, Kyoto Encyclopedia of Genes and Genomes (KEGG) pathway enrichment identified amino acid biosynthesis as one of the most significantly perturbed pathways following ETSI exposure (Fig. 1e). Gene Set Enrichment Analysis (GSEA) further confirmed significant positive enrichment for the amino acids gene set, with an enrichment score (ES) of 0.593 and normalized enrichment score (NES) of 2.680 (nominal *P* < 0.001, FDR *q* < 0.001). The positive NES indicates coordinated transcriptional up-regulation of amino acid biosynthetic machinery in ETSI treated *C. albicans* (Fig. 1f). Collectively, these transcriptomic data show that ETSI triggers broad transcriptional activation of amino acid biosynthesis pathways in *C. albicans*.

### Compound ETSI exhibits antifungal activity against *C. albicans* in an amino acid dependent manner

Given that ETSI triggers transcriptional up-regulation of amino acid biosynthetic pathways in *C. albicans*, we next sought to identify individual amino acids that could modulate its antifungal potency. Dose matrix titration assays were performed in GMM medium to assess combinatorial effects of ETSI with each of eighteen individual amino acids on *C. albicans* growth. Supplementation with most amino acids exerted minimal or negligible influence on susceptibility of *C. albicans* to ETSI (Table 1). By contrast, exogenous arginine, histidine, and lysine markedly potentiated ETSI antifungal activity, lowering its MIC from 100 μM to 6.25 μM, 6.25 μM, and 3.125 μM, respectively (Fig. 2a–c) (Table 1). To test whether this sensitization reflected a non-specific increase in fungal drug susceptibility, we combined arginine, histidine, or lysine with fluconazole. Unlike the strong potentiation observed for ETSI, none of these three amino acids produced comparable enhancement of fluconazole antifungal activity (Fig. S2a-c). These results demonstrate that the sensitization is driven by specific amino acid metabolic states, rather than a general nutritional effect of exogenous amino acids.

**Figure 2.**
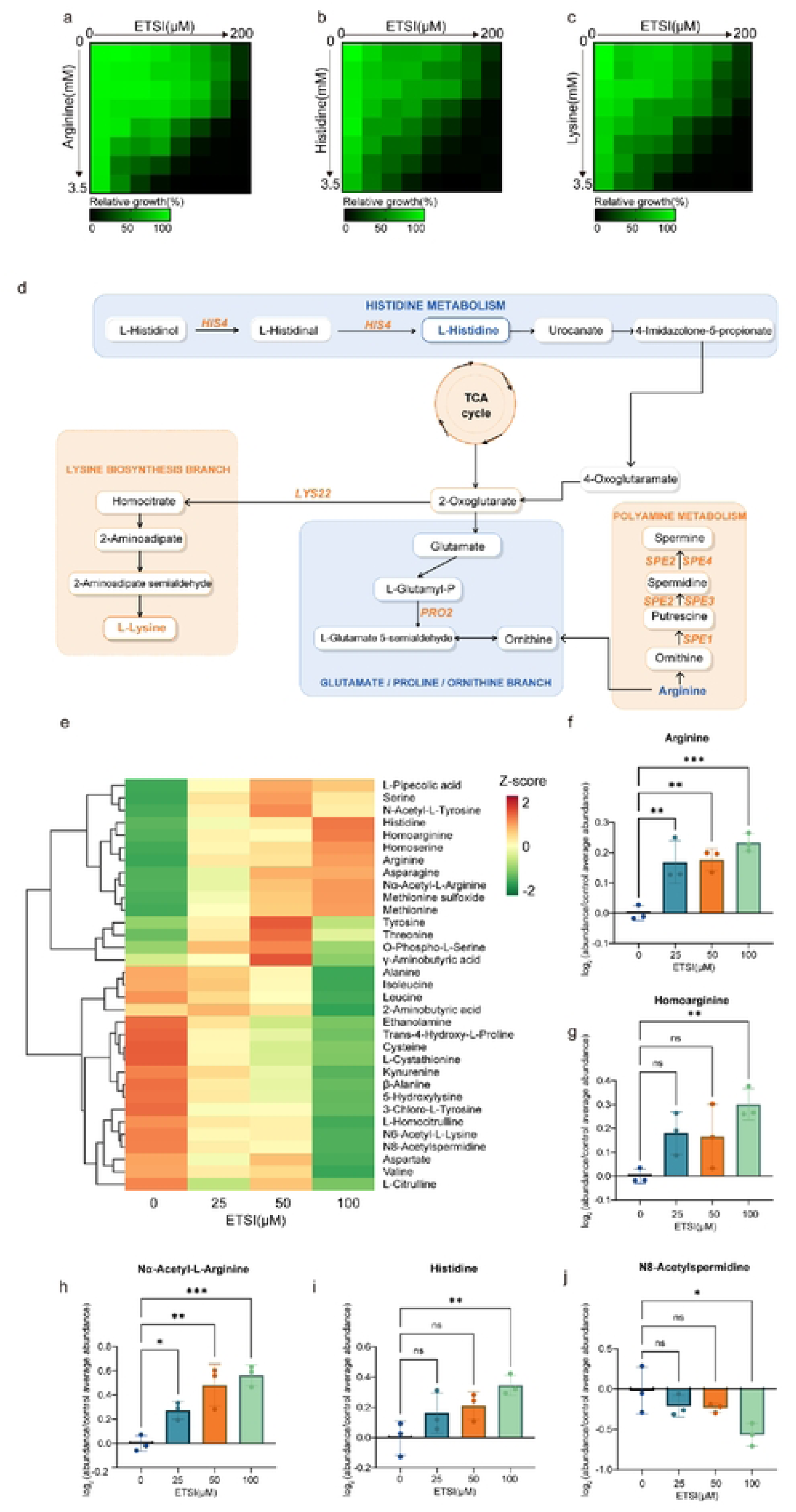
Compound ETSI exhibits antifungal activity against *C. albicans* in an amino acid-dependent manner, **a-c.** Dose-matrix titration analysis of the combination of arginine **(a),** histidine **(b)** or lysine **(c)** and compound ETSI. The color intensity quantitatively represents relative growth (as indicated by the scale bar below), **d.** Schematic representation of the metabolic relationships among arginine, histidine and lysine pathways, e. Hierarchical clustering heatmap of 33 metabolites that showed significant changes in at least one compound ETSI-treated group. Metabolite abundances were log_2_-transformcd and standardized by row (Z-score). Columns represent the mean metabolite abundance of the 0, 25, 50, and 100 μM compound ETSI groups. Red and green indicate relatively higher and lower abundance, respectively, **f-j.** Relative intracellular abundances of arginine **(f),** homoarginine (g), Nα-acetyl-L-arginine **(h),** histidine **(i),** and N8-acetylspermidine **(j)** in *C. albicans* following treatment with 0, 25, 50, or 100 μM ETSI. Metabolite abundances were normalized to the mean abundance of the untreated control and expressed as log_2_-transformed relative abundance. Data are presented as mean ± SD from three biological replicates. Statistical significance was assessed by one-way ANOVA followed by Dunnett’s multiple comparisons test (ns, not significant; ρ-value: * < 0.05, ** <0.01, *** < 0.001).

**Table 1.** Dose-matrix titration analysis of the combination of 18 amino acids with compound ETSI.

| No. | Amino Acid (AA) | MIC |  |  |
|---|---|---|---|---|
|  |  | AA(Combination) (μM) | ETSI(Alone) (μM) | ETSI(Combination) (μM) |
| 1 | Lysine | 218.8 | 100 | 3.125 |
| 2 | Arginine | 218.8 | 100 | 6.25 |
| 3 | Histidine | 218.8 | 100 | 6.25 |
| 4 | Adenine | 3500 | 100 | 50 |
| 5 | Methionine | 875 | 100 | 50 |
| 6 | Serine | 1750 | 100 | 50 |
| 7 | Threonine | 218.8 | 100 | 50 |
| 8 | Cysteine | 250 | 100 | 100 |
| 9 | Glutamic acid | 3500 | 100 | 100 |
| 10 | Leucine | 3500 | 100 | 100 |
| 11 | Phenylalanine | 1750 | 100 | 100 |
| 12 | Tryptophan | 3500 | 100 | 100 |
| 13 | Tyrosine | 3500 | 100 | 100 |
| 14 | Uridine | 3500 | 100 | 100 |
| 15 | Valine | 3500 | 100 | 100 |
| 16 | Aspartic acid | 109.4 | 100 | 200 |
| 17 | Isoleucine | 3500 | 100 | 200 |
| 18 | Proline | 3500 | 100 | 200 |

The three sensitizing amino acids, arginine, histidine, and lysine, are all basic amino acids, which share partially overlapping permease import systems including the high-affinity Can1 transporter[30], and can activate Ssy1-Ptr3-Ssy5 dependent nitrogen sensing responses in *C. albicans*[31]. While histidine and lysine also conferred sensitization to ETSI, they feed indirectly into central nitrogen pools via 2-oxoglutarate (Fig. 2d). By contrast, arginine occupies a uniquely central position in fungal nitrogen metabolism, as its catabolism directly yields ornithine, the proximal substrate for polyamine biosynthesis via Spe1-mediated decarboxylation[15](Fig. 2d). Moreover, arginine serves as a key substrate for the ammonia-generating pathway implicated in *C. albicans* escape from macrophage phagosomes. These distinctive metabolic connections linking arginine catabolism, polyamine biosynthesis, and ammonia production prompted us to further investigate whether compound ETSI perturbs intracellular arginine homeostasis and its associated metabolic pathways.

We further performed targeted amino acid metabolomic profiling of *C. albicans*. This targeted panel quantified 71 metabolites, of which 67 yielded quantifiable values in at least one treatment group. Metabolite abundances were expressed as the mean log₂-transformed relative abundance from three independent biological replicates per group (Table S2). Thirty-three metabolites showed significant changes in at least one ETSI treatment group. Hierarchical clustering of these 33 differentially altered metabolites revealed distinct concentration-dependent metabolic response patterns (Fig. 2e). Notably, arginine and several arginine related metabolites, including homoarginine and Nα-acetyl-arginine, clustered within a metabolite module with elevated levels upon ETSI exposure. Among the quantified individual metabolites, arginine exhibited the most consistent response to ETSI, with significant elevations detected at 25, 50, and 100 μM (Fig. 2f). This sustained intracellular arginine accumulation was accompanied by increased homoarginine, which reached statistical significance at 100 μM (Fig. 2g), as well as marked concentration-dependent buildup of Nα-acetyl-arginine, which rose significantly starting at 25 μM (Fig. 2h). Histidine, another basic amino acid that potentiates ETSI activity, also trended upward and was significantly enriched at the highest tested ETSI concentration (Fig. 2i). In contrast, N8-acetylspermidine, a polyamine metabolite connected to the broader arginine-ornithine-polyamine network, decreased progressively and was significantly depleted at 100 μM ETSI (Fig. 2j).

Together, these findings demonstrate that the antifungal activity of compound ETSI is strongly influenced by specific amino acids, particularly arginine, histidine, and lysine. The pronounced accumulation of intracellular arginine and its related metabolites, together with alterations in a polyamine-associated metabolite, suggests that compound ETSI disrupts arginine-associated metabolic homeostasis. Given the central role of arginine metabolism in polyamine biosynthesis and ammonia production, we therefore focused on the arginine–polyamine metabolic axis to further investigate the antifungal mechanism of compound ETSI.

### Compound ETSI attenuates *C. albicans* hyphal growth and immune evasion

Macrophages harbour an arginine rich niche, with arginine concentrations ranging from approximately 100 μM to 1000 μM[32–34]. Since this concentration range substantially potentiates ETSI antifungal activity in vitro (Fig. 2a, Table 1), we hypothesized that ETSI could exploit the arginine rich intracellular environment within macrophages to impair *C. albicans* immune evasion (Fig. 3a).

**Figure 3.**
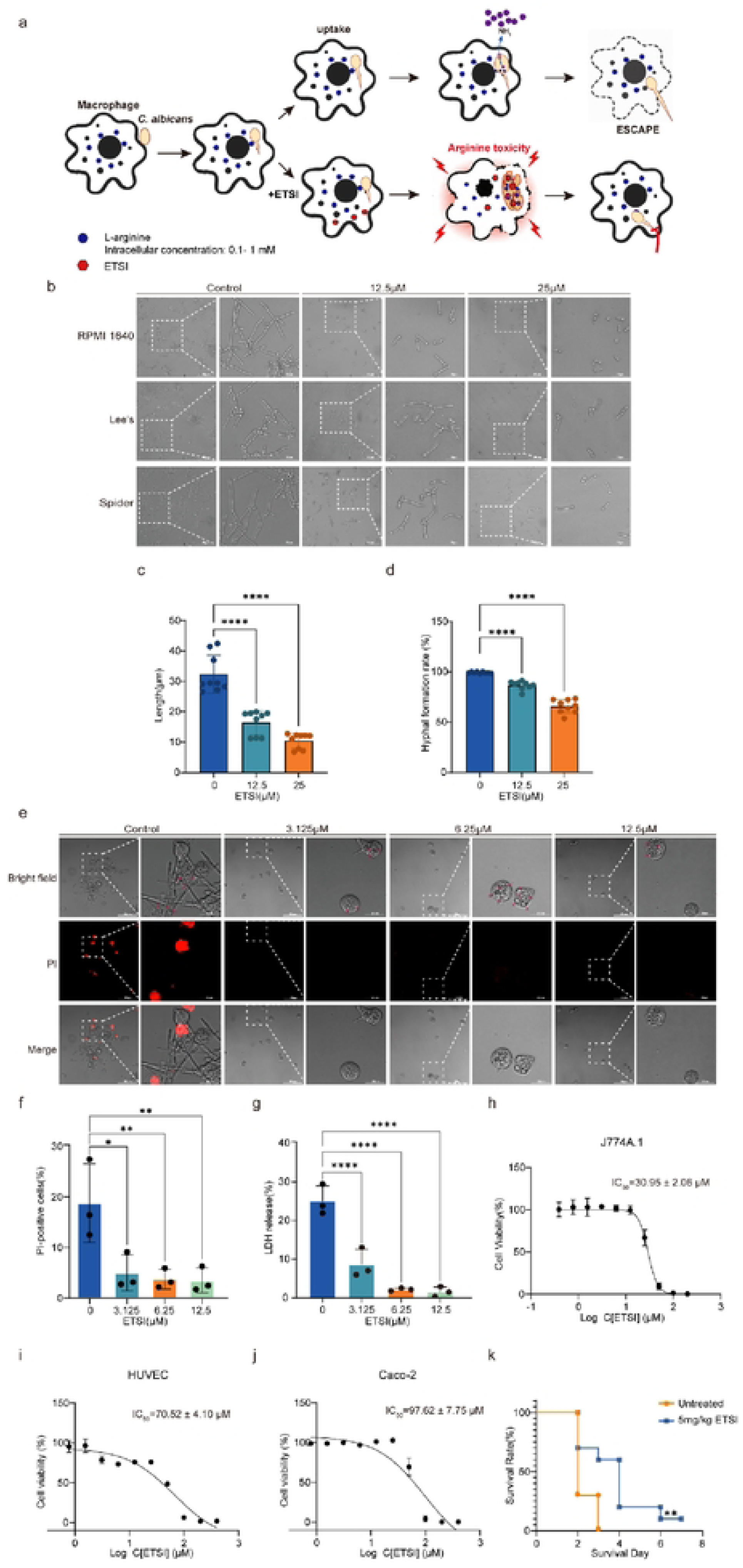
Compound ETSI attenuates hypha-associated virulence and immune evasion of *C. albicans*, **a.** Schematic illustration of the mechanism by which compound ETSI utilizes intracellular arginine in macrophages to inhibit immune evasion of *C. albicans,* **b.** Hyphal growth of *C. albicans* was induced in RPMI 1640, Lee’s, and Spider media and treated with 12.5 μM and 25 μM ETSI. Enlarged views of the boxed regions are shown adjacent to the corresponding images. Scale bars, 40 pm for overview images and 15 pm for enlarged images. Images are representative of three independent biological replicates, c. Quantification of hyphal length of *C. albicans* treated with the indicated concentrations of compound ETSI in RPMI 1640, Lee’s and Spider media. Significance was determined by one-way ANOVA with Dunnett’s multiple comparisons test, p-value: ****<0.0001. d. Quantification of the hyphal formation rate of *C. albicans* treated with the indicated concentrations of compound ETSI in RPMI 1640, Lee’s and Spider media. Significance was determined by one-way ANOVA with Dunnett’s multiple comparisons test. ρ-value: ****<0.0001. e. J774A.1 macrophages and *C. albicans* SC5314 were co-incubated at a multiplicity of infection (MOI) of 5 and treated with different concentrations of compound ETSI. After incubation, cells were stained with 20 μg/mL propidium iodide (PI). Enlarged views of the boxed regions are shown adjacent to the corresponding images. Scale bars, 80 μm for overview images and 20 μm for enlarged images. Representative images from three independent biological replicates are shown (n = 3). f. Quantification of the percentage of Pl-positive macrophages in each treatment group using ImageJ software. Significance was determined by one-way ANOVA with Dunnett’s multiple comparisons test. ρ-value: * < 0.05, **<0.01. g. J774A.1 macrophages and *C. albicans* SC5314 were co-incubated at a multiplicity of infection (MOI) of 1 and treated with different concentrations of compound ETSI. Cell death was assessed by measuring the relative release of lactate dehydrogenase (LDH) into the culture supernatant. LDH release was expressed as a percentage of the total LDH activity contributed by both the culture medium and cells. Data arc presented as mean ± SD from three independent biological experiments. Significance was determined by one-way ANOVA with Dunnett’s multiple comparisons test, p-value: ****<0.0001. **h, i, j.** Cytotoxicity of compound ETSI against J774A.l(h), HUVEC (i) and Caco-2 (j) cells assessed by the CCK-8 assay. Cell viability was determined following exposure to increasing concentrations of compound ETSI, and IC_50_ values were calculated by nonlinear regression. IC_50_ values are presented as mean ± SD from three independent experiments, k. Kaplan-Meier survival curves of *C. albicans*-infected *Galleria mellonella* treated with 5 mg/kg compound ETSI or left untreated. Survival was monitored for 7 days.p-value: **<0.01.

We first evaluated the effect of ETSI on hyphal development under multiple hypha induction conditions, including RPMI 1640, Lee’s and Spider media. ETSI markedly inhibited hypha formation at subinhibitory concentrations (Fig. 3b). In the absence of ETSI, the mean hyphal length was 32.35 ± 6.23 μm, which decreased to 16.48 ± 3.97 μm and 10.49 ± 2.50 μm following treatment with 12.5 and 25 μM ETSI, respectively, corresponding to reductions of approximately 49.0% and 67.6% relative to the untreated control (*P* < 0.0001, Fig. 3c). Consistently, the hyphal formation rate decreased from 99.18 ± 0.65% in the untreated group to 86.65 ± 4.04% and 65.70 ± 6.74% at 12.5 and 25 μM ETSI, respectively (*P* < 0.0001, Fig. 3d).

We further established a *C. albicans*-macrophage co-culture model to probe intracellular antifungal activity of ETSI. Propidium iodide (PI) staining, which evaluates macrophage plasma membrane integrity, uses a membrane-impermeable fluorescent dye excluded from intact cells but capable of penetrating compromised membranes to bind nucleic acids and emit red fluorescence[35]. Representative images show PI-positive damaged macrophages in untreated controls and *C. albicans* trapped within macrophages upon ETSI treatment (Fig. 3e). In untreated infected cultures, 18.67 ± 7.75% of macrophages were PI-positive; this fraction dropped to 4.86 ± 3.58% following 3.125 μM ETSI treatment (*P* = 0.0149). Further reductions to 3.62 ± 1.96% and 3.38 ± 2.46% were observed at 6.25 and 12.5 μM ETSI, respectively (*P* = 0.0094 and 0.0086, respectively) (Fig. 3f). We next performed lactate dehydrogenase (LDH) release assays to independently quantify macrophage damage during fungal infection (Fig. 3g). LDH is a cytosolic enzyme released upon plasma membrane rupture, so supernatant LDH levels provide a quantitative measure of host cell death. Compared with untreated infected controls showing 25.05 ± 3.92% LDH release, 3.125 μM ETSI lowered LDH release to 8.74 ± 3.81% (*P* < 0.001). Even greater protection was achieved at 6.25 μM and 12.5 μM ETSI, with LDH release decreasing to 2.18 ± 0.40% and 1.52 ± 1.35%, respectively (*P* < 0.0001). Taken together, these results corroborate that ETSI treatment protects macrophages from *C. albicans* mediated cytotoxicity.

Importantly, ETSI alone did not induce substantial PI positivity in J774A.1 macrophages at the experimental concentrations (Fig. S3a). Consistent with this observation, ETSI yielded a half maximal inhibitory concentration (IC₅₀) of 30.95 ± 2.06 μM against J774A.1 macrophages (Fig. 3h). We further evaluated mammalian cell toxicity using human umbilical vein endothelial cells (HUVEC) and human colon adenocarcinoma Caco-2 cells via Cell Counting Kit-8 (CCK-8) assays. ETSI exhibited IC_50_ values of 70.52 ± 4.10 μM and 97.62 ± 7.75 μM for HUVEC and Caco-2 cells, respectively (Fig. 3i-j). Taken together, these data reveal that ETSI exhibits limited cytotoxicity toward mammalian host cells.

To assess in vivo protective efficacy, we employed a *Galleria mellonella* larval infection model. Infected larvae received either vehicle or 5 mg/kg ETSI. ETSI treatment significantly prolonged larval survival relative to vehicle treated infected controls (Fig. 3k; *P* < 0.01). All control larvae succumbed to infection by day 3, whereas approximately 60% of ETSI-treated larvae remained alive at this time-point. Although survival gradually declined in the ETSI group, roughly 10 % of larvae were still living on day 7 post-infection. These observations show that ETSI improves host survival in an in-vivo infection setting.

### Compound ETSI antifungal activity relies on polyamine biosynthesis

Polyamine biosynthesis is tightly coupled to arginine metabolism. Within this pathway, ornithine is converted to putrescine by the ornithine decarboxylase Spe1. Putrescine is sequentially metabolized into spermidine and spermine via Spe3 and Spe4, respectively. Spe2 catalyzes the conversion of S-adenosylmethionine (SAM) to dcSAM, which provides the aminopropyl group donor required for spermidine and spermine production (Fig. 4a). Polyamines are important physiological metabolites downstream of arginine, yet excessive intracellular polyamine accumulation can trigger cytotoxicity. Notably, *SPE2*, a key gene of polyamine biosynthesis, was identified among ETSI-responsive differentially expressed genes and showed consistent transcriptional changes upon ETSI exposure (Fig. S4a). We therefore focused our further investigation on the arginine-downstream polyamine biosynthetic pathway.

**Figure 4.**
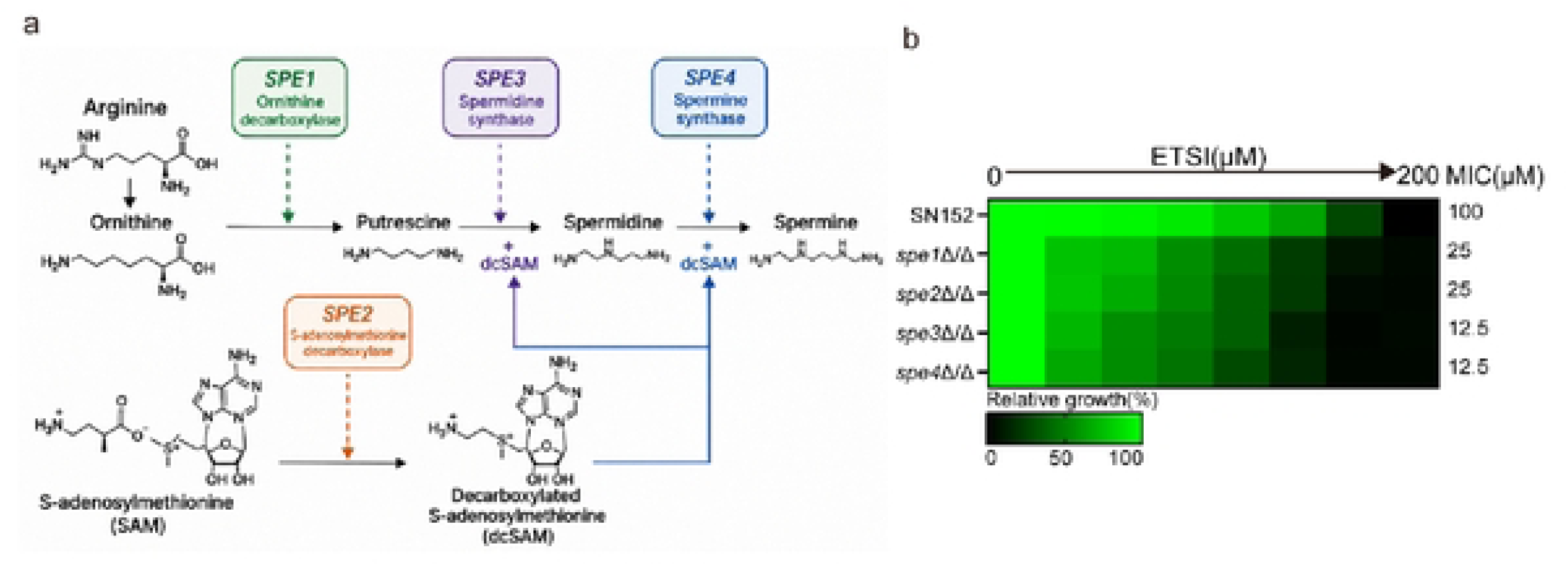
Polyamine biosynthesis pathway mediates the antifungal potentiation effect of compound ETSI. **a.** Schematic representation of the polyamine biosynthetic pathway, b. MIC values of compound ETSI against SN152, *spelΔ/Δ, spe2Δ/Δ, spe3Δ/Δ*, and *spe4Δ/Δ* were determined using the broth microdilution method in GMM medium. Color intensity quantitatively represents relative growth (sec scale bar at the bottom). The MIC of compound ETSI was defined as the lowest compound concentration resulting in more than 50% growth inhibition.

Given that arginine potentiates the antifungal activity of ETSI against *C. albicans*, we next tested whether downstream polyamine metabolites could similarly modulate ETSI potency. Exogenous spermine or spermidine markedly enhanced ETSI antifungal activity against multiple *Candida* species, including *C. albicans*, *Candida glabrata*, *Candida krusei*, *Candida parapsilosi*s, and *Candida tropicalis*, lowering ETSI MIC values from 50-200 μM to 0.391-6.25 μM (Table 2). These findings highlight the strong synergistic interactions between ETSI and either spermine or spermidine.

**Table 2.**
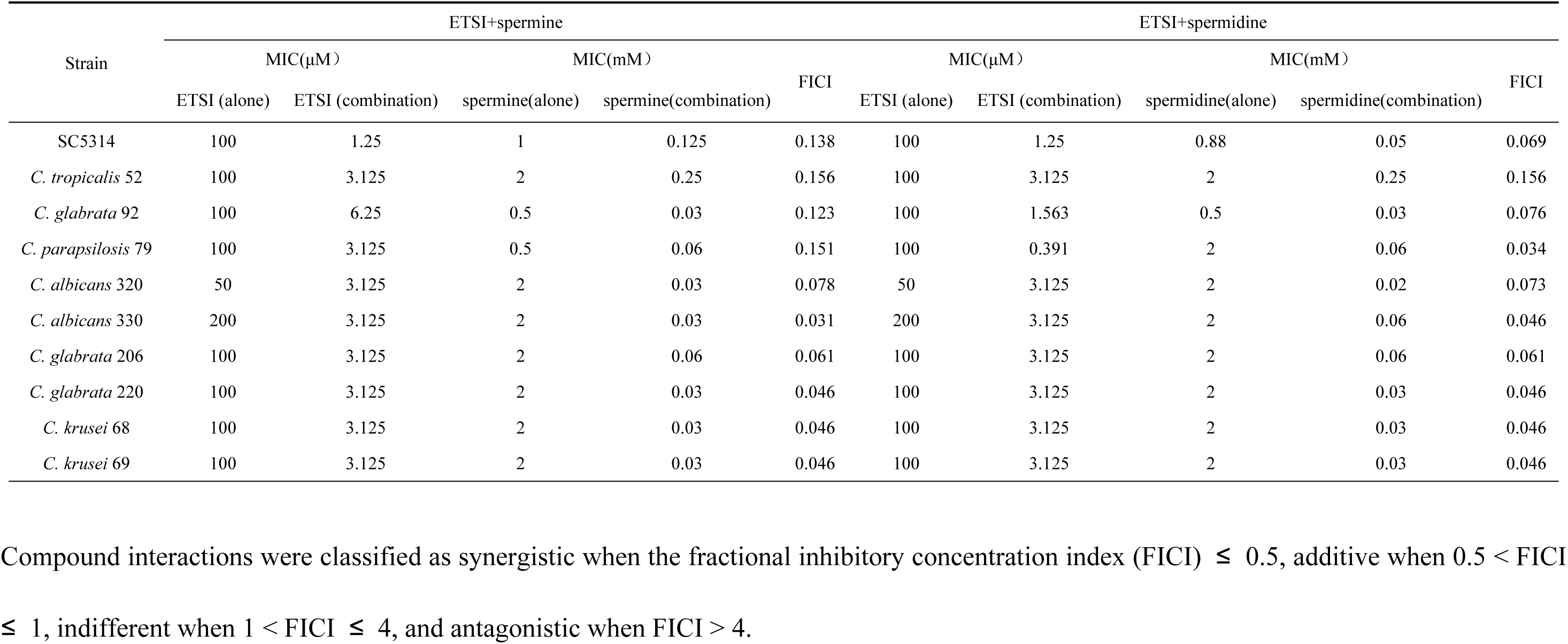
Changes in the MIC values of compound ETSI in combination with spermine or spermidine against multiple *Candida* species, including *Candida albicans*, *Candida glabrata*, *Candida krusei*, *Candida parapsilosis*, and *Candida tropicalis*.

To genetically interrogate the contribution of polyamine biosynthesis to ETSI susceptibility, we therefore generated *spe1*Δ/Δ, *spe2*Δ/Δ, *spe3*Δ/Δ, and *spe4*Δ/Δ deletion mutants (Fig. S5a). Deletion of any of these *SPE* genes rendered cells more susceptible to ETSI. The parental SN152 strain exhibited an MIC of 100 μM, whereas mutant MIC values dropped to 12.5-25 μM (Fig. 4b). When ETSI was combined with exogenous spermine or spermidine, the MIC values of ETSI against these *SPE* deletion mutants further decreased to 3.125–6.25 μM (Table 3). Of note, the magnitude of polyamine driven potentiation differed markedly between the wild-type and mutant backgrounds. In SN152, spermine reduced the ETSI MIC 16-fold (from 100 μM to 6.25 μM; fractional inhibitory concentration index, FICI = 0.071), while spermidine produced a 32-fold MIC reduction (from 100 μM to 3.125 μM; FICI = 0.039) (Table 3). By contrast, synergistic potentiation was substantially attenuated in the *spe1*Δ/Δ, *spe2*Δ/Δ, *spe3*Δ/Δ, and *spe4*Δ/Δ mutants: spermine and spermidine only reduced ETSI MIC by 2- to 8-fold, yielding FICI values ranging from 0.250 to 0.625. Although exogenous polyamines still enhanced ETSI activity in mutant strains, the diminished synergy upon loss of endogenous polyamine biosynthesis supports a functional role for this pathway in the cellular response to ETSI. Collectively, these results show that both genetic disruption of polyamine biosynthesis and exogenous polyamine supplementation lower the ETSI concentration required for robust antifungal effects, and the combination of the two perturbations produces even stronger sensitization. Intriguingly, although exogenous polyamines synergize with ETSI to suppress fungal growth, impairment of endogenous polyamine synthesis also increases ETSI susceptibility. These dual observations indicate that proper polyamine homeostasis governs *C. albicans* tolerance to ETSI imposed metabolic stress and further corroborate the involvement of the arginine-polyamine axis in shaping ETSI susceptibility.

**Table 3.** Changes in the MIC values of compound ETSI in combination with spermine or spermidine against SN152 and *SPE* deletion mutants.

| Strain | ETSI+spermine |  |  |  |  | ETSI+spermidine |  |  |  |  |
|---|---|---|---|---|---|---|---|---|---|---|
|  | MIC(μM） |  | MIC(mM） |  | FICI | MIC(μM） |  | MIC(mM） |  | FICI |
|  | ETSI (alone) | ETSI (combination) | spermine(alone) | spermine(combination) |  | ETSI (alone) | ETSI (combination) | spermidine(alone) | spermidine(combination) |  |
| SN152 | 100 | 6.25 | 2 | 0.016 | 0.071 | 100 | 3.125 | 2 | 0.016 | 0.039 |
| *spe1*Δ/Δ | 25 | 3.125 | 0.5 | 0.125 | 0.375 | 25 | 3.125 | 2 | 0.5 | 0.375 |
| *spe2*Δ/Δ | 25 | 3.125 | 0.5 | 0.25 | 0.625 | 25 | 3.125 | 2 | 0.25 | 0.250 |
| *spe3*Δ/Δ | 12.5 | 3.125 | 0.5 | 0.125 | 0.500 | 12.5 | 6.25 | 2 | 0.25 | 0.625 |
| *spe4*Δ/Δ | 12.5 | 6.25 | 2 | 0.125 | 0.563 | 12.5 | 3.125 | 2 | 0.25 | 0.375 |

Compound interactions were classified as synergistic when the fractional inhibitory concentration index (FICI) ≤ 0.5, additive when 0.5 < FICI ≤ 1, indifferent when 1 < FICI ≤ 4, and antagonistic when FICI > 4.

### Compound ETSI possibly exerts antifungal activity by disrupting V-ATPase activity

Given the functional link between polyamine homeostasis and proton gradient dependent transport, we hypothesized that ETSI activity is connected to the V-ATPase. The V-ATPase consists of a cytosolic ATP-hydrolyzing V_1_ domain and a membrane-embedded V_0_ domain, which jointly drive proton translocation across the vacuolar membrane[36, 37]. The resulting proton electrochemical gradient provides the driving force for vacuolar import of polyamines and other metabolites[38] (Fig. 5a). Impaired V-ATPase function may disrupt polyamine sequestration within the vacuole, leading to intracellular polyamine accumulation and subsequent cytotoxicity.

**Figure 5.**
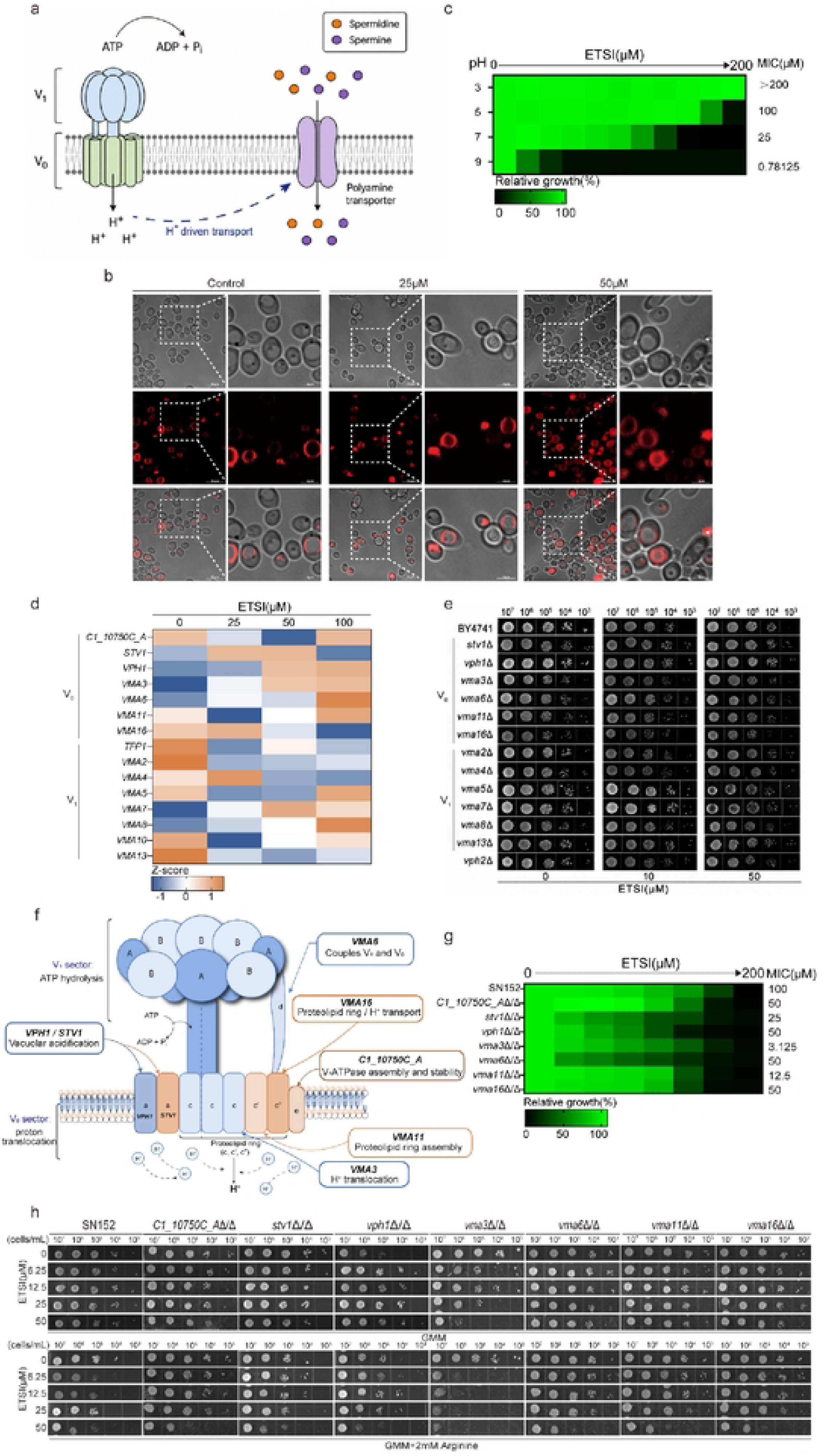
Compound ETSI may exert antifungal effects through V-ATPase inhibition, **a.** Schematic representation of the relationship between vacuolar V-ATPase and polyamine transport, b. Representative FM4-64 fluorescence images of *C. albicans* cells following treatment with 0,25, or 50 μM ETSI. Bright-field, FM4-64 fluorescence, and merged images are shown. White dashed boxes indicate the areas enlarged in the adjacent panels. Scale bars, 10 μm and 4 μm. c. Relative growth of *C. albicans* treated with different concentrations of compound ETSI (0-200 μM) at pH 3, 5, 7, and 9. The corresponding MIC values were >200, 100, 25, and 0.78125 μM, respectively, **d.** Heatmap showing the relative transcriptional expression patterns of V-ATPase-related genes in *C. albicans* following treatment with 0, 25, 50, or 100 μM ETSI. Expression values were standardized as Z-scores, with orange and blue indicating relatively higher and lower expression levels, respectively, e. Spotting assays of compound ETSI against vacuolar V-ATPase-rclatcd gene deletion mutants in YPD medium. The strains were obtained from the *Saccharomyces cerevisiae* knockout library, f. Schematic representation of the functions of the seven genes encoding the V_o_ subunits of vacuolar V-ATPase. g. MIC values of compound ETSI against *C. albicans* SN152 and seven gene deletion mutants of the V_o_ subunits of vacuolar V-ATPase were determined using the broth microdilution method in GMM medium. Color intensity quantitatively represents relative growth (see scale bar at the bottom). The MIC of compound ETSI was defined as the lowest compound concentration resulting in more than 50% growth inhibition, h. Spotting assays of compound ETSI against *C. albicans* SN152 and seven gene deletion mutants of the V_o_ subunits of vacuolar V-ATPase in the absence and presence of 2 mM arginine.

FM4-64 staining revealed marked alterations in vacuolar morphology following ETSI treatment. Untreated cells predominantly exhibited well-defined vacuolar membrane staining, whereas treatment with 25 or 50 μM ETSI resulted in increasingly irregular and diffuse FM4-64 fluorescence, indicating disruption of normal vacuolar organization (Fig. 5b). We found that ETSI exhibited a pronounced pH-dependent effect, with its MIC decreasing from >200 μM at pH 3 to 100 μM at pH 5, 25 μM at pH 7, and 0.78125 μM at pH 9 (Fig. 5c), indicating extracellular pH influences the antifungal activity of ETSI.

To test whether ETSI perturbs V-ATPase associated responses, we first profiled transcriptional changes for genes encoding V-ATPase subunits upon ETSI exposure. Multiple genes belonging to both V_0_ and V_1_ domains displayed concentration dependent expression alterations (Fig. 5d). Among V_0_ subunit associated genes, *VMA3* and *VMA6* genes were strongly up-regulated at higher ETSI concentrations, whereas the *VMA16* gene showed an overall downward trend. For the V_1_ subunit encoding genes, *VMA7* and *VMA8* genes were upregulated at higher compound ETSI concentrations; by contrast, *VMA13*, *TFP1*, and *VMA2* genes showed reduced transcript abundance (Fig. 5d).

We next performed a preliminary screen using the *Saccharomyces cerevisiae* whole genome deletion collection to survey ETSI susceptibility among V-ATPase-related mutants. Spotting assays revealed heterogeneous ETSI susceptibility across different V-ATPase deletion mutants (Fig. 5e). In mutants for V_0_-domain associated mutants, *vma3*Δ and *vma16*Δ exhibited increased susceptibility to ETSI, whereas *stv1*Δ and *vph1*Δ showed relative tolerance. Deletion of *VMA6*, or *VMA11* resulted in minor increased ETSI susceptibility. Within V_1_-domain mutants, *vma5*Δ, *vma7*Δ, and *vma13*Δ retained comparatively robust growth under ETSI treatment, while *vma2*Δ, *vma4*Δ, and *vma8*Δ displayed phenotypes similar to the parental BY4741 strain. Additionally, loss of the *VPH2* gene, encoding a V-ATPase assembly factor rather than a structural subunit of V_0_ or V_1_, led to minimal change in ETSI susceptibility (Fig. 5e). To further contextualize these findings, we mapped the functions of the V-ATPase-associated subunits within the V₁V₀ complex (Fig. 5f). These subunits contribute to distinct processes, including ATP hydrolysis, proton translocation, proteolipid ring assembly, and V₁–V₀ coupling, providing a structural framework for interpreting the differential compound ETSI susceptibility observed among V-ATPase mutants. Following this preliminary screening in budding yeast, we assessed the phenotypic consequences of deleting orthologous V-ATPase encoding genes in *C. albicans* (Fig. S5a, Fig. 5g). Individual deletion mutants for V-ATPase related genes displayed altered susceptibility to ETSI (Fig. 5g). Compared with the parental SN152 strain, mutants including *vma3*Δ/Δ, *vma6*Δ/Δ, *vma11*Δ/Δ, *vma16*Δ/Δ, *stv1*Δ/Δ, *vph1*Δ/Δ, and *C1_10750C_A*Δ/Δ all showed increased sensitivity toward ETSI, albeit to varying extents. Of these strains, *vma3*Δ/Δ exhibited the most pronounced sensitization, with an MIC reduced to 3.125 μM; *vma11*Δ/Δ and *stv1*Δ/Δ had MIC values of 12.5 μM and 25 μM, respectively (Fig. 5g).

Considering that arginine and downstream polyamines potentiate the antifungal activity of ETSI, we further tested whether exogenous arginine modulates ETSI susceptibility in these V-ATPase gene deleted *C. albicans* mutants using spotting assays (Fig. 5h). In GMM agar medium, the SN152 strain toleranted to ETSI, with no marked growth inhibition observed even at 50 μM. Deletion of V-ATPase related genes conferred variable degrees of ETSI hypersensitivity; consistent with MIC test results, the *vma3*Δ/Δ exhibited the most pronounced sensitization phenotype, showing clear growth defects at 12.5 μM ETSI. Supplementation with 2 mM arginine further enhanced the inhibitory effect of ETSI on the SN152 strain, lowering the inhibitory concentration to 6.25 μM. Arginine driven sensitization was further augmented in several V-ATPase deletion mutants, again most evidently in *vma3*Δ/Δ, in which near complete suppression was observed at 6.25 μM ETSI (Fig. 5h).

Furthermore, addition of spermine or spermidine also potentiated ETSI activity against V-ATPase gene deleted *C. albicans* mutants (Table 4). Individual deletion of these V-ATPase related genes rendered *C. albicans* more susceptible to ETSI and spermine or spermidine as single agents. However, compared with wild-type SN152, most V-ATPase mutant strains exhibited elevated FICI values for ETSI combined with spermine or spermidine, indicating weakened synergistic interactions. Notably, *vma3*Δ/Δ showed the highest intrinsic susceptibility to ETSI monotherapy (MIC = 3.125 μM, versus 100 μM in SN152). Vma3 serves as the core proteolipid subunit of the V_0_ domain, and loss of the *VMA3* gene nearly abolishes V-ATPase assembly and proton pumping activity. In this mutant, the V-ATPase complex is already functionally disrupted; hence, co-addition of spermine or spermidine failed to generate strong synergy, yielding FICI values of 0.628 for both polyamine combinations, corresponding to additive rather than synergistic effects. In comparison, other V-ATPase mutants retained partial residual proton transport activity and still exhibited some degree of synergism, though weaker than SN152 (Table 4). Collectively, although loss of V-ATPase related genes increased basal susceptibility to ETSI and polyamines, the reduced synergism upon combination treatment supports that the antifungal mechanism of ETSI is associated with functional V-ATPase.

**Table 4.** Changes in the MIC values of compound ETSI in combination with spermine or spermidine against SN152 and seven gene deletion mutants of the V_0_ subunits of vacuolar V-ATPase.

| Strain | ETSI+spermine |  |  |  |  | ETSI+spermidine |  |  |  |  |
|---|---|---|---|---|---|---|---|---|---|---|
|  | MIC(μM） |  | MIC(mM） |  | FICI | MIC(μM） |  | MIC(mM） |  | FICI |
|  | ETSI (alone) | ETSI (combination) | spermine(alone) | spermine(combination) |  | ETSI (alone) | ETSI (combination) | spermidine(alone) | spermidine(combination) |  |
| SN152 | 100 | 6.25 | 2 | 0.016 | 0.071 | 100 | 3.125 | 2 | 0.016 | 0.039 |
| *C1_10750C_AΔ/Δ* | 50 | 3.125 | 0.5 | 0.063 | 0.189 | 50 | 3.125 | 0.5 | 0.125 | 0.313 |
| *stv1*Δ/Δ | 25 | 3.125 | 2 | 0.5 | 0.375 | 25 | 3.125 | 2 | 0.5 | 0.375 |
| *vph1*Δ/Δ | 50 | 3.125 | 2 | 0.063 | 0.094 | 50 | 3.125 | 2 | 0.063 | 0.094 |
| *vma3*Δ/Δ | 3.125 | 1.5625 | 0.125 | 0.016 | 0.628 | 3.125 | 1.5625 | 0.125 | 0.016 | 0.628 |
| *vma6*Δ/Δ | 50 | 12.5 | 0.25 | 0.016 | 0.314 | 50 | 25 | 0.125 | 0.016 | 0.628 |
| *vma11*Δ/Δ | 12.5 | 6.25 | 2 | 0.25 | 0.625 | 12.5 | 3.125 | 2 | 0.25 | 0.375 |
| *vma16*Δ/Δ | 50 | 25 | 2 | 0.5 | 0.750 | 50 | 25 | 2 | 0.5 | 0.750 |

Compound interactions were classified as synergistic when the fractional inhibitory concentration index (FICI) ≤ 0.5, additive when 0.5 < FICI ≤ 1, indifferent when 1 < FICI ≤ 4, and antagonistic when FICI > 4.

### Compound ETSI targets Vma13 in *C. albicans*

To identify potential binding targets of ETSI in *C. albicans*, we performed drug affinity responsive target stability (DARTS) assay. This method relies on the principle that binding of a small molecule ligand stabilizes the conformation of its target protein, rendering it more resistant to proteolytic degradation (Fig. 6a). Total *C. albicans* protein lysates were incubated with graded concentrations of ETSI and then subjected to proteolysis with thermolysin. SDS-PAGE revealed an apparent ETSI dependent protective signal within the 40–55 kDa molecular weight range relative to the DMSO vehicle control (Fig. 6b). This gel region was excised and analysed by LC–MS/MS to identify candidate ETSI binding proteins. Guided by our preceding phenotypic findings, Vma13, a 53.3-kDa subunit of the V-ATPase, was prioritized as a candidate binding protein (Fig. 6c and Table S3).

**Figure 6.**
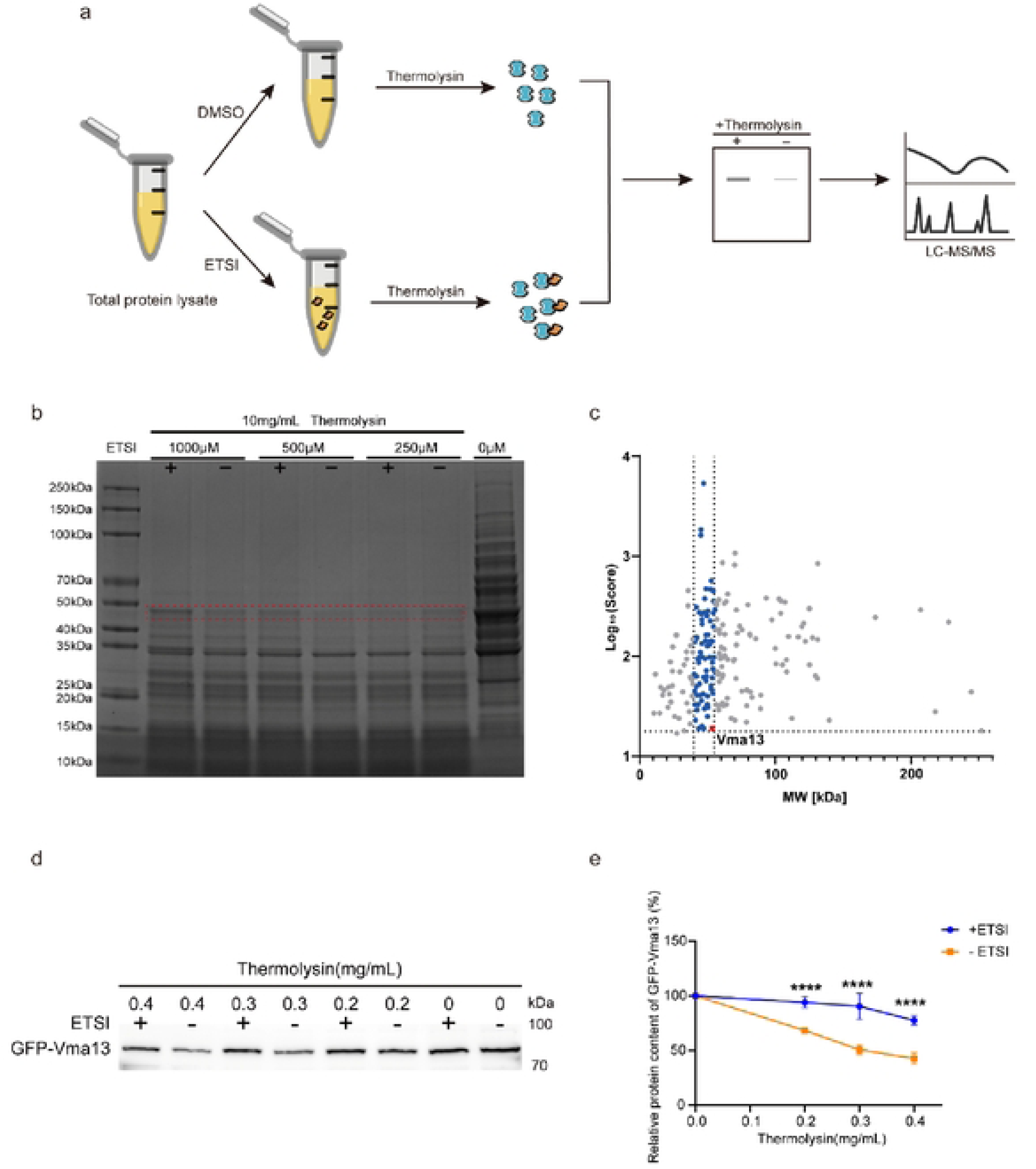
Compound ETSI targets Vmal3 in *C. albicans*. **a.** Schematic representation of the drug affinity responsive target stability (DARTS) assay used to identify potential compound ETSI-binding proteins. Total protein lysates from SC5314 were incubated with compound ETSI or DMSO, followed by limited proteolysis with thermolysin. Protein bands exhibiting compound ETSI-dependent protection from proteolysis were excised and subjected to LC-MS/MS analysis, b. Representative SDS-PAGE analysis of *C. albicans* protein lysates subjected to DARTS in the presence of 250, 500, or 1000 μM compound ETSI, followed by limited proteolysis with thermolysin (10 mg/mL). Samples treated with compound ETSI are indicated by “+”, whereas untreated samples arc indicated by “-”. The red dashed box indicates the region showing compound ETSI-dependent protection from proteolysis; the corresponding gel region was excised for LC-MS/MS analysis. Representative results from three independent biological experiments are shown, c. LC-MS/MS identification of candidate compound ETSI-interacting proteins. Candidate proteins are plotted according to molecular weight (MW) and logio-transformed identification scores, with Vmal3 specifically highlighted, d. Representative DARTS-wcstern blot validation of the interaction between compound ETSI and Vma 13. GFP-tagged Vma 13 was incubated with or without compound ETSI, followed by treatment with increasing concentrations of thermolysin (0, 0.2, 0.3, and 0.4 mg/mL), and GFP-Vmal3 was subsequently detected by immunoblotting. Representative results from three independent biological experiments are shown, e. Quantification of the relative GFP-Vmal3 band intensity following treatment with different concentrations of thermolysin in the presence or absence of compound ETSI. Protein levels were normalized to the corresponding undigested controls. Error bars represent the standard deviations of three independent biological experiments (n = 3). *p* -value: ****<0.0001.

To further verify this interaction, in *C. albicans*, we constructed a GFP-tagged Vma13 mutant strain and performed DARTS followed by western blot analysis. At each thermolysin concentration tested, the GFP-Vma13 band intensity was consistently greater in ETSI-treated samples than in matched DMSO controls, indicating that ETSI increased Vma13 resistance to proteolysis. While GFP-Vma13 abundance declined progressively in both groups as thermolysin concentration rose, higher levels of GFP-Vma13 were preserved in samples supplemented with ETSI (Fig. 6d). Densitometric quantification of immunoblotting results validated this protective effect: at thermolysin concentrations of 0.2-0.4 mg/mL, the residual GFP-Vma13 level was significantly higher in ETSI-treated samples compared with controls (*P* < 0.0001) (Fig. 6e). Collectively, these DARTS results show that ETSI protects Vma13 from proteolytic digestion, supporting Vma13 as a candidate binding target of ETSI in *C. albicans*.

## Discussion

Here, we functionally characterized ETSI, a thiadiazole-succinimide derivative, as a unique arginine-dependent antifungal compound. While ETSI shows minimal direct growth inhibition under arginine free conditions, it displays robust activity against *C. albicans* within arginine rich microenvironments mimicking the macrophage intracellular niche. Mechanistically, ETSI targets Vma13 to suppress V-ATPase H⁺ pumping activity. V-ATPase inhibition disrupts vacuolar H⁺ sequestration and causes cytoplasmic proton overload in *C. albicans*, which drives two critical antifungal consequences. On one hand, excess intracellular H⁺ converts diffusible NH_3_ into membrane impermeable NH_4_^+^, sequestering NH_3_ within fungal cells and abolishing phagosomal alkalinization. On the other hand, collapse of the V-ATPase H⁺ electrochemical gradient impairs vacuolar metabolite storage, resulting in cytoplasmic accumulation of arginine and polyamine metabolites. Accumulated polyamines trigger profound metabolic cytotoxicity. Combined, these dual metabolic disruptions block yeast-to-hypha transition, lock *C. albicans* in the yeast morphology, and disable fungal immune escape from macrophages (Fig. 7). These results reveal a previously undescribed mechanism connecting V-ATPase H⁺ balance to NH_3_ efflux and polyamine toxicity, offering a niche-targeted therapeutic paradigm against invasive candidiasis.

**Figure 7.**
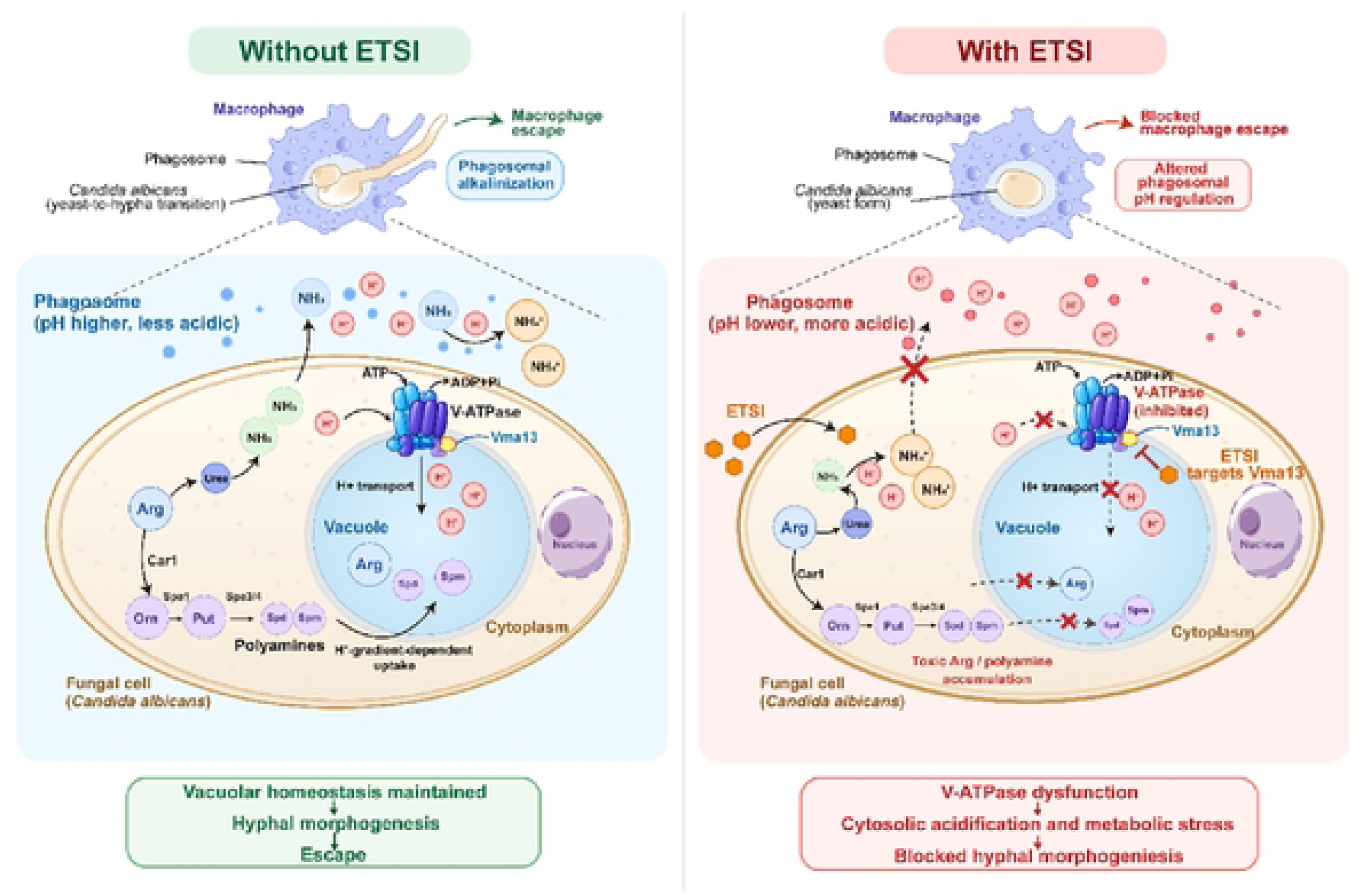
Proposed model for ETSI-mediated blockade of *C. albicans* macrophage escape through disruption of Vmal3/V-ATPase-dependent arginine-polyamine homeostasis. In the absence of ETSI, vacuolar V-ATPase maintains the proton electrochemical gradient by pumping cytoplasmic H^+^ into the vacuole, thereby supporting vacuolar acidification and sequestration of arginine and polyamine metabolites. Arginine catabolism bifurcates downstream of Carl: urea is further degraded by Durl,2 to generate NH_3_ and CO_2_, whereas ornithine enters the polyamine biosynthetic pathway through Spel and subsequently Spe3/Spe4. These two branches respectively contribute to phagosomal pH modulation and polyamine homeostasis. In the presence of ETSI, interaction with the V_r_H subunit Vmal3 impairs V-ATPase activity and proton translocation, resulting in cytoplasmic H^+^ retention and disruption of the vacuolar proton gradient. Increased cytoplasmic proton availability shifts intracellular ammonia toward NH_4_* and limits NH_3_ release, contributing to altered phagosomal pH regulation. Meanwhile, impaired proton-gradient-dependent vacuolar sequestration causes cytoplasmic accumulation of arginine and disturbances in polyamine metabolism, leading to metabolic stress and cytotoxicity. Together, these perturbations suppress yeast-to-hypha morphogenesis, retain *C. albicans* predominantly in the yeast form, and block macrophage escape.

Macrophage phagocytosed *C. albicans* deploys multiple strategies to evade killing, including morphological switching from yeast to hyphae, neutralization of phagosomal acidification, reactive oxygen species (ROS) detoxification and secretion of virulent hydrolases. Among these adaptive programs, arginine metabolism serves as a critical metabolic hub coupling nutrient acquisition, pH remodeling and hyphal morphogenesis to support intracellular survival and subsequent escape. The fungal Stp2-mediated arginine uptake pathway fuels dual metabolic cascades: ammonia production for phagosomal alkalinization and hyphal morphogenesis, and polyamine biosynthesis essential for fungal physiological homeostasis. Under physiological conditions, vacuolar V-ATPase uses ATP hydrolysis in the cytosolic V₁ sector to drive proton translocation through the membrane-embedded V₀ sector, thereby establishing a stable proton electrochemical gradient[39]. This gradient provides the driving force for proton-gradient-dependent vacuolar sequestration of arginine, polyamines and other metabolites, thereby preventing their excessive accumulation in the cytoplasm. Importantly, Vma13 is not merely a structural component of the V-ATPase complex. Studies in *S. cerevisiae* have shown that the V₁-H subunit encoded by the *VMA13* gene contributes to coupling ATP hydrolysis to proton translocation, such that reduced Vma13 function can preserve substantial ATPase activity while markedly impairing generation of the vacuolar proton gradient[40, 41].Our mechanistic results demonstrate that ETSI interacts with Vma13 and suppresses V-ATPase activity, which dissipates vacuolar H⁺ gradients and abolishes the vacuolar compartmentalization capacity for arginine and polyamine metabolites. Direct biochemical studies in isolated yeast vacuoles have demonstrated that spermidine and spermine uptake is ATP dependent and requires the H⁺ potential generated by vacuolar H⁺-ATPase; disruption of V-ATPase activity markedly inhibits this process[38]. Likewise, arginine transport into isolated yeast vacuoles requires ATP and functional V-ATPase activity[42]. Although these direct transport measurements were primarily performed in *S. cerevisiae*, the conserved role of V-ATPase in fungal vacuolar acidification provides a strong mechanistic basis for proposing similar H⁺ gradient dependent sequestration in *C. albicans*. Integrated metabolomic profiling combined with genetic and chemical rescue experiments further validates that aberrant accumulation of arginine serves as the primary cause of fungal metabolic cytotoxicity. This metabolic toxicity impairs fungal ammonia generation and phagosomal neutralization, locks *C. albicans* in the yeast form, and ultimately abrogates macrophage immune escape, establishing a complete causal linkage between Vma13/V-ATPase dysfunction, polyamine metabolic disturbance, and fungal immune evasion arrest.

Vma13/V-ATPase dysfunction may additionally influence ammonia handling through altered H⁺ homeostasis. Reduced H⁺ translocation into the vacuole would be expected to favor cytoplasmic H⁺ retention. Because NH₃ and NH₄⁺ exist in rapid pH-dependent equilibrium (NH₃ + H⁺ ⇌ NH₄⁺), increased cytoplasmic H⁺ availability shifts the equilibrium toward protonated NH₄⁺ and reduces the fraction of freely diffusible NH₃[43]. Because NH₄⁺ is positively charged and therefore far less permeable across the hydrophobic lipid bilayer than uncharged NH₃, this shift further restricts passive ammonia diffusion across the fungal plasma membrane. Consequently, compound ETSI may reduce the freely diffusible NH₃ pool and thereby limit ammonia efflux across the fungal plasma membrane.

Notably, the requirement for phagosomal alkalinization in *C. albicans* immune escape remains debated. Several studies have reported that hyphal elongation and macrophage escape can proceed independent of phagosomal neutralization, arguing that morphological switching itself, rather than pH elevation, serves as the primary driver of escape. Our results do not contradict these observations. Rather, our data indicate that arginine metabolism contributes to dual outputs: ammonia-mediated phagosomal alkalinization and polyamine-dependent fungal fitness. In our experimental system, ETSI mediated Vma13 inhibition reduces ammonia production and impairs alkalinization, but the resultant cytotoxic accumulation of arginine-polyamine metabolites acts as an additional, pH-independent mechanism restricting intracellular survival and morphogenesis. Thus, suppression of alkalinization represents one contributing factor, while polyamine metabolic disruption constitutes the primary driver arresting fungal escape in our model.

Prior studies have well documented the critical role of V-ATPase in maintaining fungal vacuolar function, stress tolerance and virulence[44–46]. Most functional evidence originates from genetic ablation of VMA subunits or pharmacological inhibition using pan-V-ATPase inhibitors such as bafilomycin A1. However, bafilomycin A1 globally suppresses V-ATPase activity in both fungal and mammalian cells, leading to severe host cytotoxicity and limiting its translational applicability. Although genetic disruption of individual VMA subunits attenuates *C. albicans* virulence, these genetic perturbations cannot be recapitulated by small-molecule targeting, and no subunit-selective V-ATPase modulators have been reported for antifungal intervention. Consistent with our DARTS assay results, ETSI interacts with Vma13 rather than broadly inhibiting the entire V-ATPase complex, distinguishing it from these pan-inhibitors. In the context of fungal polyamine metabolism, previous studies have primarily focused on biosynthetic inhibition to reduce polyamine levels and constrain fungal proliferation. In contrast, ETSI does not interfere with polyamine biosynthesis but instead impairs vacuolar metabolite sequestration, triggering toxic accumulation of endogenous arginine metabolites, a phenotype directly supported by our metabolomic profiling and validated via genetic and chemical rescue experiments. This mode of action fundamentally differs from conventional metabolic inhibitors that suppress synthetic pathways. Moreover, most existing anti-*C. albicans* compounds block hyphal growth through direct regulation of morphogenetic signaling cascades, accompanied by robust growth suppression. ETSI, by comparison, exhibits negligible antifungal activity under arginine-depleted conditions and only functions within arginine-rich intracellular niches, uncoupling general growth inhibition from specific immune-evasion blockade. A key innovation of this study lies in the unique arginine-dependent, niche-specific antifungal activity of ETSI, consistent with our in vitro growth assays showing minimal activity in arginine-limited media and robust inhibition under arginine-replete conditions, which distinguishes it from all current clinical antifungals and experimental antifungal leads. Conventional azoles, echinocandins and polyenes target constitutive fungal structural or synthetic pathways and exert uniform growth inhibition regardless of microenvironmental metabolic status, which imposes persistent growth pressure and facilitates the emergence of drug resistance. In contrast, ETSI specifically disables the adaptive metabolic circuit that empowers intracellular *C. albicans* to survive and escape macrophage clearance. To our knowledge, ETSI is the first reported small molecule that interacts with fungal Vma13 to disrupt V-ATPase dependent arginine-polyamine homeostasis and block *C. albicans* macrophage immune evasion. Additionally, ETSI exhibits minimal cytotoxicity against host macrophages, supporting a favorable therapeutic window for intracellular antifungal therapy.

This study has several limitations. First, although DARTS assays confirm the specific interaction between ETSI and Vma13, further biophysical assays including SPR or ITC will help quantify binding affinity and precisely map the binding interface. Second, the *Galleria mellonella* model provides preliminary in vivo evidence; future validation in murine invasive candidiasis models is required to further evaluate translational potential and pharmacological performance. Third, structure activity relationship optimization of thiadiazole-succinimide derivatives is needed to improve compound potency, solubility and druggability. Despite these limitations, our current work establishes a solid mechanistic framework for targeting fungal vacuolar metabolic homeostasis and highlights Vma13 as a tractable antifungal target to block intracellular immune evasion, offering valuable insights for the development of next-generation metabolism-based antifungal therapeutics.

## Methods and materials

### Strains, compounds, cell lines, primers, and culture conditions

All fungal primers and strains used in this study are listed in Tables S4 and S5, respectively. Unless otherwise specified, *C. albicans* strains were routinely cultured in YPD medium containing 1% (w/v) yeast extract, 2% (w/v) peptone, and 2% (w/v) glucose at 30 °C with shaking. For solid culture, YPD medium was supplemented with 2% (w/v) agar.

Antifungal susceptibility assays were performed in GMM, consisting of 0.67% (w/v) yeast nitrogen base (YNB), 0.5% (w/v) ammonium sulfate, and 2% (w/v) glucose. Where indicated, GMM was supplemented with the corresponding amino acids at the concentrations specified for each experiment. For hyphal induction assays, *C. albicans*cells were cultured in RPMI 1640 medium buffered to pH 7.0 with MOPS, Lee’s medium, or Spider medium, as indicated for the respective experiments.

All compounds used in this study were dissolved in dimethyl sulfoxide (DMSO; Sangon Biotech, Shanghai, China) and stored at −20°C until use. Where appropriate, vehicle controls containing equivalent concentrations of DMSO were included.

Human colorectal adenocarcinoma cells (Caco-2; YCL-0363), human umbilical vein endothelial cells (HUVECs), and murine monocyte/macrophage cells (J774A.1) were obtained from the American Type Culture Collection (ATCC; HTB-37). Cells were maintained in high-glucose Dulbecco’s modified Eagle’s medium (DMEM) supplemented with 10% fetal bovine serum (FBS) and 1% penicillin–streptomycin (100×; CellorLab, Shanghai, China) at 37°C in a humidified atmosphere containing 5% CO₂.

*Saccharomyces cerevisiae* deletion mutants were obtained from a commercially available yeast deletion library (Open Biosystems, YSC1053, USA) and routinely cultured in YPD medium at 30°C. *Candida albicans* deletion mutants, including *SPE*- and V-ATPase-related mutants, were generated by PCR-mediated homologous recombination using auxotrophic selection markers. Transformants were selected on appropriate synthetic defined media lacking the corresponding nutritional supplements according to the selection marker used, and correct gene replacement was verified by diagnostic PCR using the primers listed in Table S4.

### Antifungal susceptibility assay

Antifungal susceptibility testing was performed according to the Clinical and Laboratory Standards Institute (M27-A3), with a few modifications[47]. First, 100 μL of 400 μM of the drug was serially diluted on a 96-well plate, and 100 μL of *C. albicans* at a concentration of 1 × 10^3^ cells/mL was added. The plate was incubated at 30°C for 24 h. The optical density at 600 nm (OD₆₀₀) of each well was measured using a Multiskan Sky microplate reader. The lowest concentration of the compound that inhibited 50% of fungal growth compared with the control was defined as the MIC value. The same procedure was applied to the remaining strains.

Dose matrix titration assay was performed essentially as described for MIC determination[28, 48]. Drug A and drug B were serially twofold diluted in two separate 96-well plates along the row and column directions, respectively. Then, 50 μL from each corresponding well of the two plates was transferred into the same well of a third 96-well plate. Subsequently, 100 μL of a *C. albicans* suspension (1 × 10³ cells/mL) was added to each well, and 200 μL of medium without drugs was added to column 9 as a negative control. The plate was incubated at 30°C for 24 h, and the optical density at 600 nm (OD₆₀₀) was measured using a Multiskan Sky microplate reader.

Dose matrix titration assays were conducted to evaluate the combinational effects of the compound with the following 18 amino acids: uridine, adenine, L-arginine, L-aspartic acid, L-glutamic acid, L-histidine, L-leucine, L-lysine, L-methionine, L-phenylalanine, L-serine, L-threonine, L-isoleucine, L-tryptophan, L-tyrosine, L-valine, L-cysteine, and L-proline. Each amino acid was used at an initial concentration of 3.5 mM, and the compound was used at an initial concentration of 200 μM. Both agents were twofold serially diluted and combined in 96-well microtiter plates for treatment against *C. albicans*. The GMM culture medium consisted of YNB supplemented with glucose and ammonium sulfate. After incubation at 30°C for 24 h, fungal growth was assessed by measuring the optical density at 600 nm (OD₆₀₀) using a Multiskan Sky microplate reader.

### RNA sequencing

RNA-seq analysis was conducted with minor modifications to published methods[49, 50]. *C. albicans* was incubated at 30 °C for 16 hours. The overnight cultures were diluted 1:100 into fresh YPD medium, resulting in a final concentration of approximately 1 × 10⁶ cells/mL. The diluted cultures were then divided into four groups (with three biological replicates per group) and treated with compounds at concentrations of 25μM (1/2 × MIC), 50μM (1×MIC), and 100μM (2×MIC), with an equivalent volume of DMSO used for the control group. All samples were cultured at 30 °C with shaking for 8 hours. After treatment, the cultures were centrifuged to collect the cell pellets, which were then sent to BGI Genomics (Shenzhen, China) for eukaryotic transcriptome re-sequencing. Differentially expressed genes were analyzed using KEGG pathway enrichment and Gene Ontology (GO) enrichment analyses on their website. Differentially expressed genes (DEGs) were defined as those with an absolute log_2_ fold change greater than 1 and a Q value below 0.001.

### Hyphal growth assay

The hyphal growth assays were conducted following the previously described method[9, 50]. *C. albicans* was cultured overnight in YPD at 30°C for 16 h. They were washed twice with PBS and resuspended in RPMI 1640 medium to a final concentration of 2 × 10⁵ cells/mL. In a 24-well plate, 500 µL of cell suspension with or without different concentrations of the compound was added to each well and incubated at 37°C for 4 h. Hyphal morphology was observed using a standard inverted microscope (Motic AE2000, China). For quantitative analysis, hyphal lengths were measured using ImageJ software, and the hyphal formation rate was calculated as the percentage of cells exhibiting hyphal growth relative to the total number of cells counted. Data obtained from RPMI 1640, Lee’s medium, and Spider medium were pooled for the quantitative analysis of hyphal length and hyphal formation rate.

### CCK-8 cytotoxicity assay

CCK-8 cytotoxicity assay was conducted according to previously established methods[50]. Human umbilical vein endothelial cells (HUVECs), human colon adenocarcinoma Caco-2 cells and macrophage cells (J774A.1) were seeded into 96-well plates at a density of 1 × 10⁵ cells/mL (100 µL per well) and incubated overnight at 37°C and 5% CO₂. After removal of the culture medium, 100 µL of two-fold serially diluted compounds were added to wells in columns 1–11, while 100 µL of medium was added to column 12 as the negative control. Cells were incubated for 24 h at 37°C and 5% CO₂. Subsequently, 10 µL of CCK-8 reagent was added to each well, followed by further incubation for 2 h at 37°C in 5% CO₂. Absorbance was then measured at 450 nm using a Multiskan Sky microplate reader.

### Lactate dehydrogenase (LDH) cytotoxicity assay

LDH release assay was performed in accordance with the previously established protocol[10]. J774A.1 cells were seeded into a 96-well plate at a density of 2 × 10⁵ cells/mL, with 100 μL of cell suspension added to each well. The cells were cultured in DMEM medium containing 10% FBS and 1% penicillin-streptomycin at 37°C and 5% CO₂ for 24 h. The culture supernatant was then aspirated, and 200 μL of 2 × 10⁵ cells/mL *C. albicans* cell suspension and different concentrations of compound ETSI (3.125 μM, 6.25 μM and12.5 μM) was added to each well; an untreated control group was also established. After 5 h of co-incubation with *C. albicans*, the culture supernatant was collected, and LDH activity was measured according to the instructions of the Lactate Dehydrogenase (LDH) Cytotoxicity Assay Kit (Beyotime, Shanghai, China) to evaluate the effect of compound ETSI treatment on the extent of damage to J774A.1 cells caused by *C. albicans*.

### Spotting assay

The spotting assay was conducted in accordance with a previous protocol[28, 51]. The overnight-cultured strains were first adjusted to a concentration of 1×10⁷ cells/mL using sterile phosphate-buffered saline (PBS). A tenfold serial dilution was then performed to prepare five concentration gradients, and 3 μL samples were spotted onto solid culture plates containing different concentrations of the compound. The plates were then incubated at 30°C for 48 h, after which the growth of the strains was recorded using an automated chemiluminescence imaging analyzer (Tanon, Shanghai, China).

### Construction of gene deletion mutants

*C. albicans* gene deletion mutants were constructed using a PCR-mediated homologous recombination strategy[52]. For each target gene, the 5′ and 3′ flanking regions were amplified from *C. albicans* SN152 genomic DNA using gene-specific primers. The selectable marker, *HIS1* or *ARG4*, was amplified separately. The 5′ flank, selectable marker and 3′ flank were then fused by fusion PCR using primers P1 and P6 to generate gene deletion cassettes for subsequent transformation. The *HIS1*-containing PCR product was first transformed into *C. albicans* cells to replace one allele of the target gene through homologous recombination. Positive transformants were selected on the corresponding selective medium. The second allele was disrupted in the same manner using the *ARG4*-containing PCR product. Mutants in which both alleles of the target gene were successfully replaced were used for subsequent experiments.

### DARTS

The DARTS assay was conducted following a previously established protocol with minor modifications[50, 53, 54]. Firstly, the total protein of *C. albicans* SC5314 was extracted. The cells washed with PBS were suspended in Detergent-free Lysis buffer (5mM EDTA, 1mM PMSF, 1X protease inhibitor), and the glass beads were mixed with the resuspended bacterial solution in a grinding tube and shaken 4 times for 30s each time. The total protein solution of the supernatant obtained by centrifugation was diluted to 5mg/mL and then co-incubated with compound ETSI or DMSO for 1 hour at 37℃. The samples were then enzymatically digested with thermolysin for 20 min, respectively, and the digestion was terminated with 0.5M EDTA. 5×Loading buffer was added for SDS-PAGE and Coomassie brilliant blue staining (Yamei, Shanghai, China) was used to determine the bands that specifically bound to ETSI. Bands were cut out and subjected to LC-MS after trypsin digestion. The peptide mixtures were separated by liquid chromatography and then analyzed using tandem mass spectrometry. Peptides were analyzed by LC–MS using a Thermo LTQ-Orbitrap mass spectrometer coupled to an Eksigent LC pump. The raw MS data were searched against the corresponding protein database to identify candidate proteins.

To evaluate the interaction between ETSI and Vma13 by DARTS, total protein lysates were prepared from the GFP-Vma13 strain as described above. Aliquots of the lysates were treated with 1000 μM ETSI or the corresponding vehicle control and incubated for 1 h at 37 ℃. The samples were then subjected to limited proteolysis with thermolysin at 0.2, 0.3, or 0.4 mg/mL for 15 min, with an enzyme-free sample included as the undigested control. Following proteolysis, the protein samples were resolved by 4–20% SDS-PAGE and electrotransferred onto polyvinylidene difluoride (PVDF) membranes. GFP-Vma13 was detected by immunoblotting with an anti-GFP primary antibody, followed by an HRP-conjugated secondary antibody. Chemiluminescent signals were visualized using the Pierce ECL detection system (Thermo Fisher Scientific, USA).

### Laser confocal microscopy

For PI staining of macrophage cell line J774A.1, 2 × 10⁵ cells/mL J774A.1 cells were seeded in confocal culture dishes and incubated overnight at 37°C under 5% CO₂. J774A.1 cells were then cocultured with a mixture of 1 × 10^6^ cells/mL *C. albicans* and different concentrations of compound ETSI for 6 hours and stained with 20 μg/mL PI (MCE, Shanghai, China) for 15 minutes. Observations were performed using a Zeiss laser scanning confocal microscope (LSM 900, Carl Zeiss AG, Oberkochen, Germany) in which PI fluorescence was excited at 561 nm.

To detect the effect of compound ETSI on vacuole morphology, *C. albicans* SC5314 was adjusted to 1 × 10^7^ cells/mL, and treated with different concentrations of compound ETSI or DMSO for 8 hours. The cells were collected and added to FM4-64 (Med Chem Express, HY-103466) for staining. The cells were incubated with shaking for 30min. The plates were then resuspended in 500uL YPD medium and incubated for 30min with continued shaking. Finally, the supernatant was discarded by centrifugation, resuspended in 100uL of YPD, and FM4-64 fluorescence was visualized using a Zeiss confocal laser scanning microscope equipped with a 63× oil-immersion objective.

### *Galleria mellonella* survival analysis

*Galleria mellonella* larvae were randomly assigned to two groups (n = 10 per group): PBS control and ETSI (5 mg/kg). All larvae were infected with *C. albicans* SC5314 at an inoculum of 7 × 10^5^ cells per larva. The fungal suspension was administered by injection in a volume of 10 μL per larva. At 2 h post-infection, larvae received a single 10-μL injection of vehicle or the indicated treatment. Following treatment, the larvae were maintained at 30 °C, and survival was monitored daily for 7 days. Larvae were considered dead when they showed no response to physical stimulation.

### Targeted amino acid metabolomics analysis

*C. albicans* SC5314 was cultured in YPD medium at 30 °C for 16 h. Overnight cultures were diluted 1:100 into fresh YPD medium to a final cell density of approximately 1×10^6^ cells/mL. The cultures were subsequently divided into four groups, with three independent biological replicates per group. Cells were treated with ETSI at 25 μM (1/2×MIC), 50 μM (1× MIC), or 100 μM (2× MIC) whereas the control group received an equivalent volume of DMSO. All cultures were incubated at 30 °C with shaking for 8 h. Following treatment, cells were harvested by centrifugation, the culture supernatants were discarded, and the resulting cell pellets were collected for targeted amino acid metabolomic analysis. Samples were submitted to Metware Biotechnology Co., Ltd. for metabolite extraction and liquid chromatography–tandem mass spectrometry (LC–MS/MS) analysis.

Metabolite extraction, LC–MS/MS detection, data preprocessing, metabolite identification, and relative quantification were performed according to the standardized workflow of Metware Biotechnology. Metabolites were identified and quantified with reference to the Metware Database (MWDB). For downstream analysis, metabolite abundances were compared among the 0μM (control), 25μM (1/2×MIC), 50μM (1× MIC), or 100μM (2× MIC) groups. For data visualization, metabolites exhibiting significant alterations in at least one treatment group were selected for hierarchical clustering analysis. Metabolite abundance values were standardized to Z-scores for each metabolite and visualized as a heatmap. For selected metabolites associated with basic amino acid and arginine metabolism, including arginine, homoarginine, Nα-acetyl-L-arginine, histidine, and N8-acetylspermidine, relative metabolite abundances were further analyzed across the four ETSI concentrations. Statistical analyses were performed using three biological replicates per condition.

### Statistical analysis

Data are presented as mean ± standard deviation (SD) unless otherwise indicated. The number of independent biological replicates is specified in the corresponding figure legends. Statistical analyses were performed using the appropriate tests according to the experimental design. For comparisons involving multiple treatment groups, the multiple-comparison procedure used is specified in the corresponding figure legend. Survival curves were analyzed using an appropriate time-to-event comparison test. Statistical significance was defined as *P* < 0.05.

## Acknowledgements

This study received financial support from the National Natural Science Foundation of China (No.82574464 and No. 82473973).

## Data Availability Statement

The RNA-seq data generated in this study have been deposited in the NCBI Sequence Read Archive under BioProject accession number PRJNA1307052 (https://www.ncbi.nlm.nih.gov/sra/PRJNA1307052). The proteomics data generated in this study have been deposited in the iProX repository under accession number IPX0019067000 (https://www.iprox.cn/page/project.html?id=IPX0019067000). The metabolomics data generated in this study have been deposited in the MetaboLights repository under accession number MTBLS16028 (https://www.ebi.ac.uk/metabolights/MTBLS16028).

## Author Contributions

**Xinyu Fang** conducted most of the experiments and performed data analysis.**Xinyu Fang**, **Yuanying Jiang**, and **Hui Lu** wrote the manuscript draft and revised the manuscript. **Xinyu Fang**, **Bingchen Jiang**, and **Wanqian Li** constructed mutant strains. **Xinyu Fang**, **Xiaojing Xu**, **Liping Xu**, **Xiao Zhou,** and **Wenqing Shi** evaluated the antifungal activities and data collection. **Xinyu Fang**, **Yuanying Jiang**, and **Hui Lu** discussed and analyzed the data. **Yuanying Jiang** and **Hui Lu** conceived the idea.**Yuanying Jiang** and **Hui Lu** directed the experiments.

